# Utility of Occupancy Models with Environmental DNA (eDNA) from Olympic Coast National Marine Sanctuary

**DOI:** 10.1101/2021.08.04.455111

**Authors:** Christopher Paight, Jenny Waddell, Matthew P Galaska

**Affiliations:** Department of Biological Sciences, University of California Santa Barbara, Santa Barbara, CA, 93101, U.S.A.; NOAA Olympic Coast National Marine Sanctuary, Port Angeles, WA, 98362; Cooperative Institute for Climate, Ocean, & Ecosystem Studies, NOAA Pacific Marine Environmental Lab, University of Washington, Seattle, Washington, USA, Seattle, WA, 98115

## Abstract

Adjacent to the spectacular, rugged coastline bordering the Olympic Peninsula of Washington State, Olympic Coast National Marine Sanctuary (OCNMS) encompasses 8257 km^2^ of coastal waters that support one of North America’s most productive marine ecosystems. These rich waters support important recreational, commercial and subsistence fisheries for Washington state and four sovereign tribal governments: the Quinault Indian Nation and the Makah, Quileute and Hoh Tribes. Given its ecological, cultural and economic significance, managers from OCNMS are tasked with monitoring and management towards conserving the area’s ecological integrity. The development of metabarcoding with environmental DNA (eDNA) as an effective freshwater monitoring tool has potential applications in marine systems, but the statistical analysis and the interpretation of those eDNA results are still under development. One promising strategy to analyze eDNA is through the use of occupancy models. Occupancy models enable us to calculate probabilities of detection from each taxon sequenced and are designed to work with presence-absence data. To inform our long term monitoring design of this coastal ecosystem, we collected 44 eDNA samples at nine of OCNMS’ long term mooring sites in 2019 and tested four different molecular markers for species reported and taxonomic richness. Additionally, we assessed occupancy models for use with eDNA to estimate the number of samples required to accurately predict the presence of a species. Occupancy models show great promise for use in eDNA studies provided there is sufficient replication; additionally, the choice of molecular marker strongly influences a taxa’s probability of detection.

**Author Summary:** We show the utility of using occupancy models with eDNA in Olympic Coast National Marine Sanctuary for monitoring near-shore marine biological communities. Based on probabilities of detection, long term coastal monitoring projects using eDNA would benefit from designs prioritizing sample number (10-20 samples per site) over sites sampled. With proper study design, occupancy models provide a statistical framework for comparisons between sites and over time. By accounting for simple non-detection vs true absence, occupancy models help to eliminate noise from eDNA studies, increasing detection of true shifts in community composition. We also demonstrate that marker choice is an important study design consideration not only for the types of taxa recovered, but also for consistency between samples and timepoints.

## Introduction

Bordered by the scenic coastline of Washington’s Olympic Peninsula and the temperate rainforest and wilderness coastline of Olympic National Park, lies Olympic Coast National Marine Sanctuary (OCNMS) (Figure 1). Designated in 1994, OCNMS protects 8257 km^2^ of coastal waters along the Olympic Peninsula of Washington State. The sanctuary encompasses a rich upwelling zone, which supports one of America’s most productive fisheries [1]. The cold, high nutrient waters delivered to the continental shelf through seasonal (April to October) upwelling support Washington’s largest and most persistent kelp beds [2]and provide essential habitat for federally protected species including Northern Sea Otters (*Enhydra lutris*), Tufted Puffins (*Fratercula cirrhata*), Killer whales (*Orcinus orca*), and humpback whales (*Megaptera novaeangliae*). Sanctuary managers are tasked with conducting research, assessments, and monitoring to support ecosystem based management as part of NOAA’s sanctuary management plan [3].

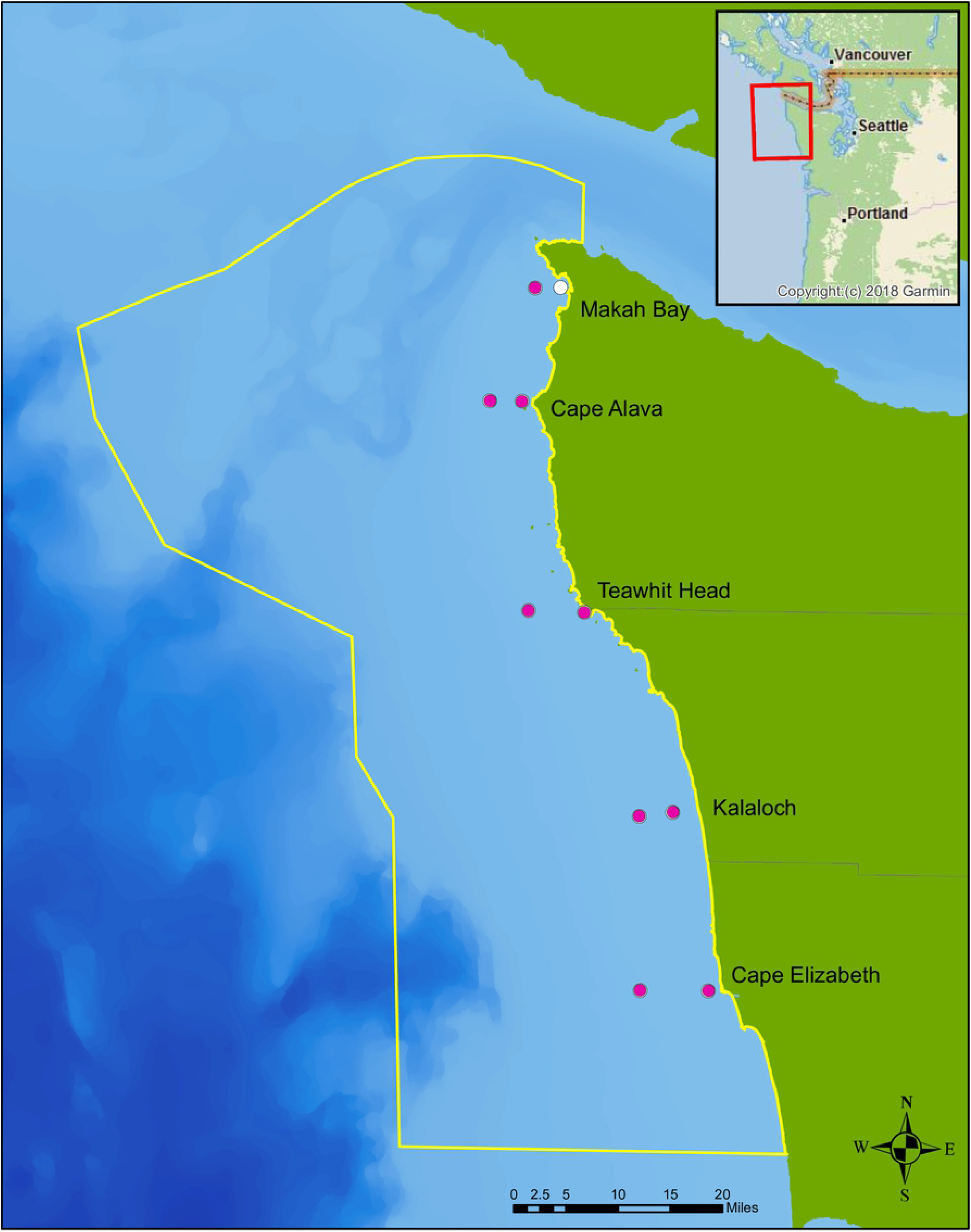

To meet these management goals, OCNMS conducts and coordinates seafloor mapping efforts, collects oceanographic measurements, explores deep sea coral and sponge communities, monitors harmful algal blooms, records underwater sound, conducts SCUBA transects through kelp forests, and performs surveys for a variety of species including intertidal communities and sea otters (“Olympic Coast National Marine Sanctuary - Science”). These efforts have generated data about the physical and biological makeup of OCNMS despite challenges that limit the types of studies, frequency of sampling that can be done, and constraints on the species that can be surveyed regularly by the sanctuary’s small staff. Ocean acidification and hypoxia, both consequences of anthropogenic climate change, are predicted to impact upwelling regions first and are a threat to the habitats that support this productive region [4].

Impacts from climate change are already being observed in the Pacific Northwest, from significant losses to oyster aquaculture caused by acidification, to major die off and displacement of fish and Dungeness crab from hypoxia [5,6]. As ocean acidification and hypoxic events are expected to increase in frequency and severity [7], it is vital to be able to understand how these changes will impact the biological community as a whole. Until recently it has been difficult to collect biological community data, particularly in remote and difficult to access habitats like OCNMS. However, over the past ten years, developments in environmental DNA (eDNA) and metabarcoding methods have made such community wide studies much more accessible [8].

There are many advantages of using metabarcoding with eDNA, but foremost among them is ease of sample collection [9]. Collecting filtered water samples integrates easily into existing research cruise operations without requiring significant additional time, equipment, or even specific sampling locations. This means that eDNA collection does not require specialized cruises, eliminating the most expensive part of any oceanographic study: ship time. Working within sanctuaries and other protected habitats often involves permits, but because eDNA sampling is non-destructive with minimal to no environmental impacts, obtaining permits for eDNA studies may be simpler. Ease of collection and storage also makes eDNA ideally suited for automation [10]. Automated collection from moorings provides finer time scale resolution than ship time allows. Automated water filtration collectors could be added to existing assets that currently collect physical and chemical oceanographic data, adding a biological component to mooring array data.

In addition to the advantages in collection, eDNA is also capable of distinguishing taxa that are morphologically too similar to discriminate, greatly increasing the numbers and types of taxa that can be monitored [11–13]. Environmental DNA studies are easy to customize, since the design and selection of markers can be tailored to fit regions and taxa of interest [14]. However, as we show in this manuscript, not all markers behave the same way, and the utility of any given marker needs to be carefully assessed. In many applications metabarcoding with eDNA has been shown to be more sensitive and comprehensive than traditional methods [15–17].

Many managers are already using eDNA/metabarcoding for environmental assessments and management decisions [18–25]. Nearly all of these monitoring programs are in freshwater systems, and the utility of eDNA/metabarcoding for coastal communities is not well known. Differences in taxonomic diversity, water exchange, and scale likely influences the detection and interpretation of eDNA, making direct transfer of freshwater eDNA methods and designs to coastal marine systems without validation inadvisable. This necessitates the development of coastal adapted sampling designs.

Although eDNA is an incredibly useful tool, there are limitations when using it for ecological assessments and monitoring [26]. The primary limitation is that metabarcoding with eDNA is not quantitative and makes comparisons between sites and time points relative [8,27]. Absolute quantification of metabarcoding data may be impossible with current methods and technologies; relative quantification between samples falls short when compared to the standards of more traditional methods [28]. Short amplification lengths and incomplete sequence representation in databases limits high confidence with respect to taxonomic assignment to species or genus with current methods. This is particularly apparent in coastal marine systems, where high rates of biodiversity in the ecological communities are often poorly represented in public databases [29,30].

One way around taxonomic uncertainty is to restrict analysis to the amplicon sequence variants (ASVs) themselves. Using ASV’s eliminates problems caused by sequence clustering, but is not without its own drawbacks [31]. An artifact of amplicon sequencing is the introduction of non-random errors which create false ASVs. These errors are termed false haplotypes, and they greatly increase the apparent diversity [32–34]. Error correction methods like those found in DADA2 help to de-noise these errors, but they are not perfect [35]. The false haplotypes add complications to downstream analysis and need to be accounted for in long term monitoring projects. Another problem is the high variation between replicates, both biological and even between technical replicates [36,37]. For long term monitoring, the high variability between samples is problematic, and proper study design needs to ensure adequate samples are collected to account for this irregularity. Metabarcoding with eDNA studies must also be able to estimate true absence from simple non-detection, which has proved challenging [26].

Limitations with eDNA need to be considered in designing a study robust enough to be replicable and sensitive enough to detect changes in sites and between years. Sufficient sampling and appropriate statistical methods are needed to account for the variability and the semi-stochastic nature of eDNA. The recent application of occupancy modeling, a common community ecology method, has the potential to address many of these critical design aspects of eDNA studies [38,39]. We incorporated occupancy models to address two of the major challenges posed by eDNA: variability between replicates and true absence. Occupancy models account for imperfect detection in presence/absence data by making the assumption that if a taxa was found in one location, it was present in that location in subsequent sampling trips, and times when a taxa was not detected were simply non-detections not absence [40–42]. With only slight modification, these methods can be adapted to work with eDNA metabarcoding data [39]. These models enable the calculation of a taxon’s probability of detection, its site occupancy, and required sample number.

In order to develop a robust, long term, ecological monitoring plan using eDNA for OCNMS we collected, processed and preserved 44 eDNA samples adjacent to nine OCNMS long term coastal moorings in summer 2019 and used occupancy models to analyze the data. We tested four different molecular markers, two of which are general markers designed to amplify a wide range of eukaryotic species, and two targeted markers designed to amplify only specific taxa--in this case metazoans and bony fish. Markers were assessed for species detected, overall richness of biodiversity, and probability of detection. With the results, we estimate required sample numbers, examine differences in molecular marker behavior, test the hypothesis that OCNMS is a contiguous biological community, and demonstrate the utility of applying occupancy models in metabarcoding with eDNA studies in coastal habitats. Finally, we explain how occupancy models with eDNA data can provide managers with statistically rigorous results upon which to base management decisions.

## Results

In total 44 1L water samples were collected from nine sites in Olympic Coast National Marine Sanctuary. One of the filters from Cape Alava 15m was damaged; this sample was discarded. Filter extraction yielded an average of 200 ng/µl of DNA with a minimum of 25 ng/µl and a maximum of 866 ng/ul. Sequence read numbers from each successive round of quality trimming are shown in the supplementary table (Supplementary table S1). An error in the COI reverse Illumina adaptor sequence prevented sequencing of COI reverse reads. All data reported for COI is from forward reads only. Error correction in DADA2 produced 894 amplicon sequence variants (ASVs) for 18S rRNA, 6159 COI, 1,020 16S rRNA, and 931 12S rRNA (Supplementary data S2). When the ASVs were taxonomically assigned, 226 taxa were recovered from 18S rRNA, 194 taxa were recovered from COI, 126 taxa were recovered from 16S rRNA, and 39 taxa were recovered from 12S rRNA (Figure 2a, Supplementary table S3). Species accumulation curves for each of the four markers indicate sampling was not exhaustive and further taxa would be recovered with more sampling (Figure 2b). A summary of the taxonomic makeup of each marker is provided in supplementary figure S4. Count tables of the number of reads for each ASV and the taxonomy for each of the four markers are found in supplementary tables (Supplementary table S5).

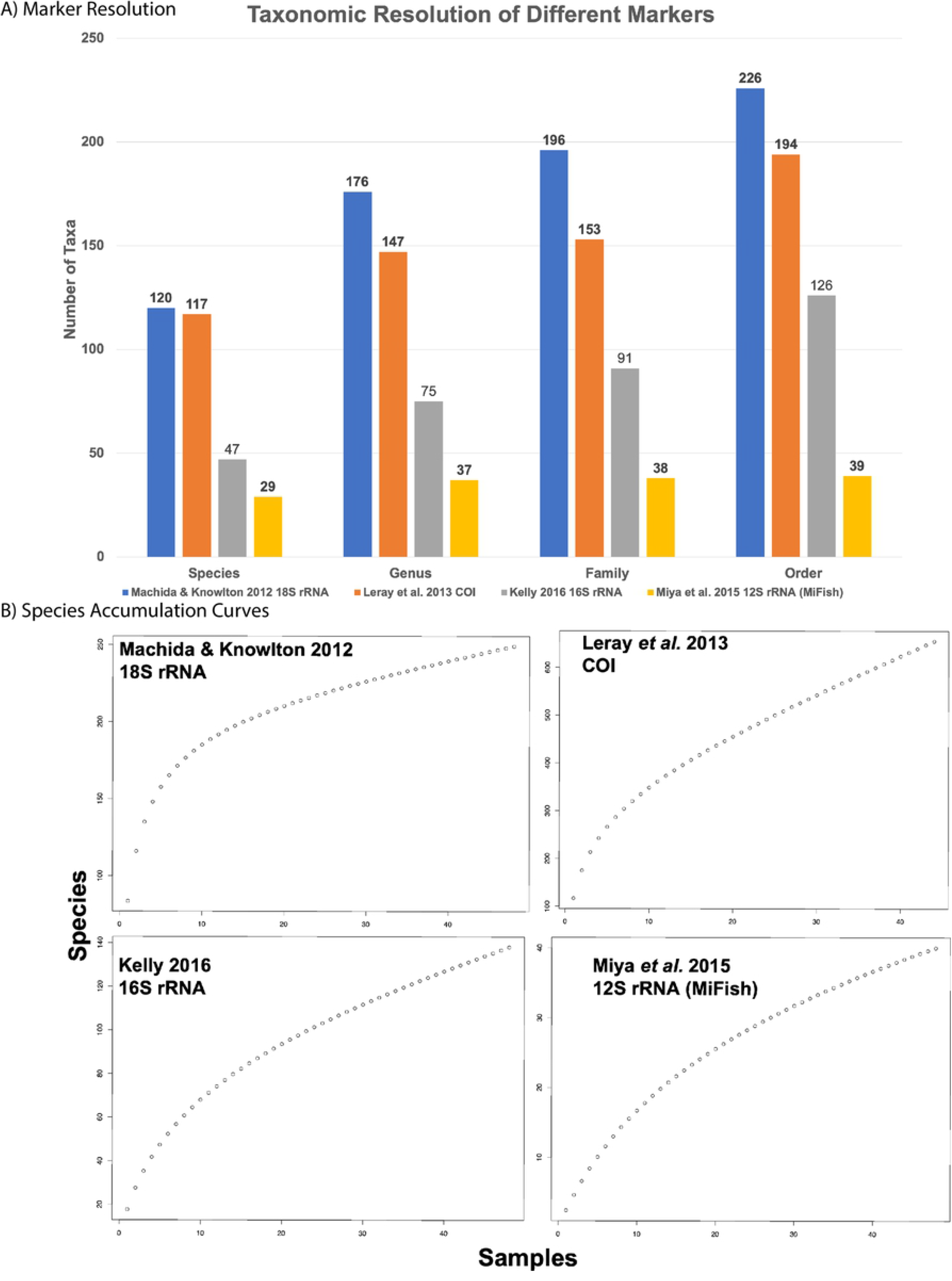

Figure 2) Taxonomy recovered and species accumulation curves for the four markers A) Taxonomic resolution of the four markers at different taxonomic ranks (species, genus, family, order). Bars show taxon number identified to each particular taxonomic rank (blue 18S, orange COI, grey 16S, yellow 12S).

B) Total species accumulation curves from the four markers. Note different y-axes.

Several taxa of note for their commercial, cultural, or special protective status were recovered by these four markers. The 16S rRNA assay recovered Killer whale (*Orcinus orca*), Gray whale (*Eschrichtius robustus*), Sea otter (*Enhydra lutris*), and Harbor porpoise (*Phocoena phocoena*). The 12S rRNA assay recovered Herring (*Clupea harengus*), Pacific cod (*Gadus macrocephalus*), Coho salmon (*Oncorhynchus kisutch*), Chinook salmon (*Oncorhynchus tshawytscha*), Steller’s sea lion (*Eumetopias jumbatus*), and California sea lion (*Zalophus califorianus*). The COI assay recovered Pacific razor clam (*Siliqua patula*). None of these taxa were specifically targeted, but all are of special interest to OCNMS and are an important target of future monitoring efforts.

### Occupancy modeling results

The majority of ASVs from all four markers have predicted site occupancy of 100%, but also have extremely large confidence intervals (Figure 3a). The low confidence in occupancy is due to the low probability of detection for these ASVs (Figure 3b). While all four markers have a large proportion of low probability of detection ASVs, there is a distinct trend of proportionally, with more low probability detections in the more specific markers 16S rRNA and 12S rRNA. Interestingly, differences between general versus specific markers extend beyond probability of detection differences. Total sequence reads, number, and probability of detection are strongly correlated for both 18S rRNA (r=0.87, R^2^=0.75) and COI (r=0.85, R^2^=0.73). However, in the more specific markers this relationship is weaker for 16S rRNA (r=0.53, R^2^=0.29) and 12S rRNA (r=0.50, R^2^=0.24) (Figure 3c). Using probabilities of detection calculated from the occupancy models we can calculate the number of samples required to have 95% confidence that an ASV is not present at a given site (Figure 3d). For the ASVs with the lowest probability of detection (0.02) 130 biological replicates are required. The steep declining asymptotic slope means higher probabilities of detection reach 95% confidence in absence with as few as 5 samples. Probabilities of detection for taxa are similar to the distributions of ASVs for each marker.

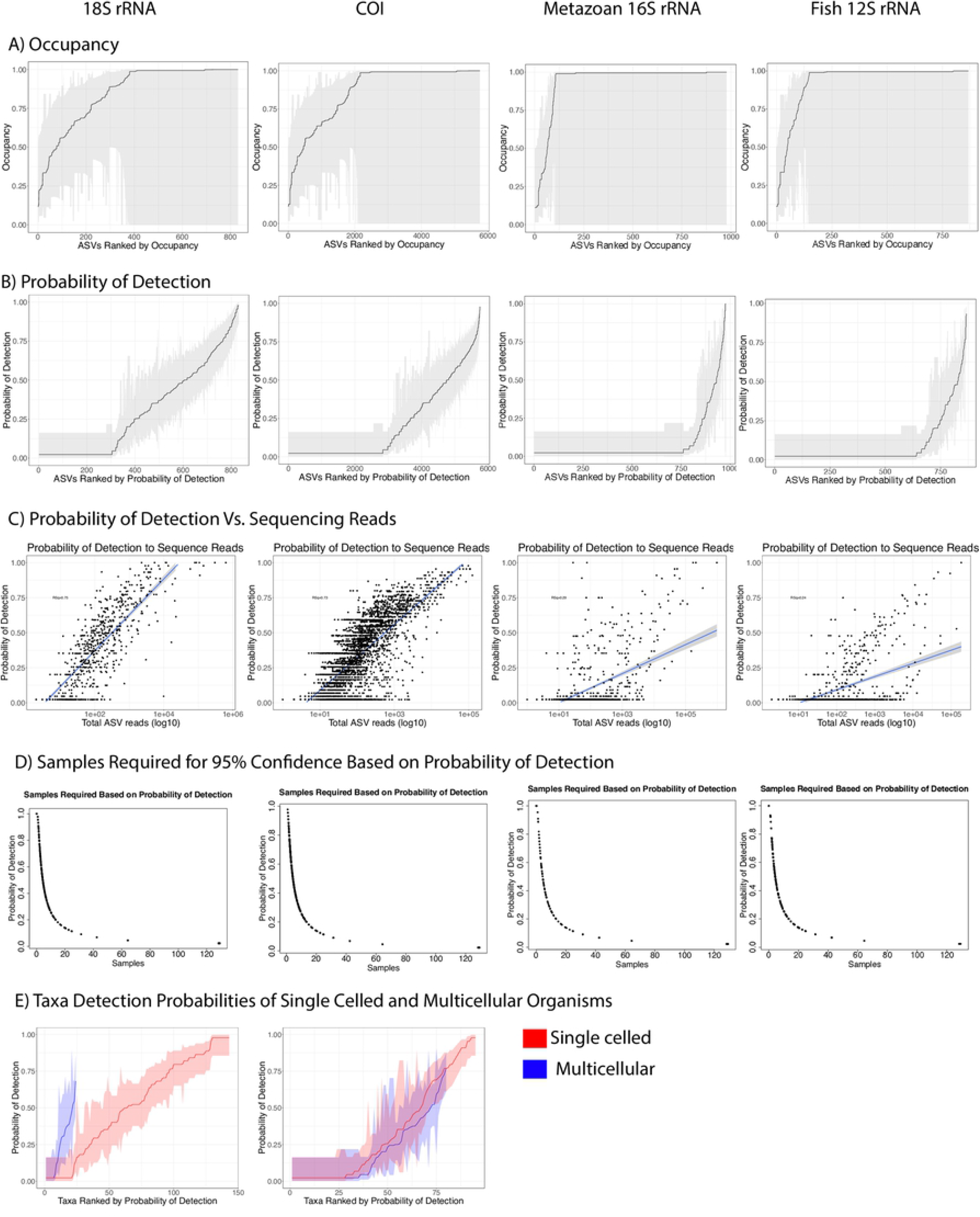

Figure 3) ASV occupancy levels and probability of detection Each of the four markers correspond to a column of plots (18S, COI, 16S, 12S) A) ASV’s ranked by occupancy level (black line) showing the 95% confidence interval (grey shading). B) ASV’s ranked by probability of detection (black line) showing the 95% confidence interval (grey shading). C) Plot of individual ASV’s total sequence reads to its probability of detection (black dots). Linear correlation in probability of detection to Log10 transformed total sequencing reads (black line) with 95% CI (blue shading). 18S rRNA (r=0.87, R2=0.75), COI (r=0.85, R2=0.73),16S rRNA (r=0.53, R2=0.29), and 12S rRNA (r=0.50, R2=0.24) D) Samples required based on an ASV’s probabilities of detection. Note ASV’s with the same probability of detection are shown as a single point on the graph. E) Taxa’s probability of detection separated by whether taxa is single-celled (red line) with 95% confidence interval (red shading) or multicellular (blue line) with 95% confidence interval (blue shading). Only 18S and COI had enough single celled/multi-celled taxa for comparison; other markers only detected multicellular organisms.

Interestingly, there are only slight differences in detection probability when single celled organisms are compared to multicellular organisms (Figure 3e). No multicellular organisms had greater than 0.80 probability of detection, but the relative proportion of probabilities is nearly identical between single celled and multicellular organisms. This is interesting given that it is still unclear whether eDNA in the ocean is primarily contained within intact cell nuclei or if it is “free floating”.

The four makers had large variation in the percentage of ASVs/taxa above the 95% confidence level that an ASV/taxa was truly absent. However, in all markers there were steeply diminishing returns above 20 samples (Table 1). With ten replicates, the 95% confidence threshold for a taxons absence is 0.26, at this level; 50% of 18S rRNA ASVs and 70% of taxa are retained, compared with just 13% ASVs and 22% of taxa for the 12S rRNA marker. With 20 replicates the probability threshold is 0.14 which includes an additional 8% (58% total) ASVs and 9% (79% total) taxa for 18S rRNA. The 12S marker had a similar increase, 6% more ASVs (19% total), and 12% more taxa (34% total). With 20 samples 79% of taxa recovered by 18S have sufficient sampling,

**Table 1).** Percentage of ASV’s/Taxa retained with different sampling numbers.

| Number of Samples | ASVs |  |  | Taxa |  |  |
| --- | --- | --- | --- | --- | --- | --- |
|  | 10 | 20 | 30 | 10 | 20 | 30 |
| Probability of Detection | 0.26 | 0.14 | 0.095 | 0.26 | 0.14 | 0.095 |
| Marker | Percentage of ASVs above detection threshold |  |  | Percentage of taxa above detection threshold |  |  |
| 18S rRNA | 50.24 | 58.33 | 60.95 | 70.06 | 79.04 | 81.44 |
| COI | 34.40 | 43.97 | 45.97 | 40.57 | 54.86 | 57.71 |
| 16S rRNA | 11.22 | 15.10 | 16.12 | 33.33 | 43.94 | 44.70 |
| 12S rRNA | 13.02 | 19.12 | 21.08 | 22.50 | 35.00 | 40.00 |

while at the same sampling level only 34% of taxa recovered by 12S do. Adding an additional 10 replicates (30 total) lowers the probability threshold to 0.095%, but only increases ASVs and taxa retained by ∼2% across all markers. Four taxa of special interest, Herring (Clupea harengus), Pacific cod (Gadus macrocephalus), Chinook salmon (Oncorhynchus tshawytscha), and Coho salmon (Oncorhynchus kisutch) have probabilities of detections above the 20 replicates threshold.

None of the ASVs or taxa detected had significantly different distributions either with a north to south divide or near shore to offshore divide (Supplementary table S6). These models were run to determine if the nine sites sampled in OCNMS were a contiguous biological community with eDNA. Based on the lack of statistical significance in the models we do not have enough data to refute that OCNMS is a contiguous biological community. With 5 replicates per site, the 95% confidence level for non-detection of an ASVs/taxa’s probability of detection needs to be greater than 0.45, limiting the number of taxa that can be used to compare specific sites. For the two targeted markers this is particularly evident do to the removal of nearly all of the taxa that would enable site to site comparisons.

## Discussion

The practical application of monitoring biological communities with eDNA/metabarcoding is already being utilized by several organizations and bio-assessment companies in terrestrial and freshwater systems. However, its utility in coastal marine systems is not well understood. Highly dynamic water exchange, greater taxonomic diversity, and size of the water body need to be considered. In designing a long-term monitoring project, sample size, number, sites, and markers need to be carefully planned to ensure the design is sensitive enough to detect changes and robust enough to be repeatable. A key component of any experimental design’s sensitivity and repeatability is sufficient replication. This is particularly true with methods that have high variability between biological replicates as well as technical replicates, as is the case with eDNA [43]. A related issue is differentiating true absence from simple non-detection [44]. Quantifying absence is essential when dealing with presence/absence data and is of particular importance for managers conducting long term ecological monitoring. Determining sufficient replication, and to an even greater degree, separating true absence from non-detection has been a major challenge for eDNA studies.

Our study aims to develop a robust sampling scheme for long term ecological monitoring of OCNMS and to provide a reference for other coastal marine monitoring projects. All four of the markers tested (18S rRNA, COI, 16S rRNA, and 12S rRNA) recovered unique taxa not detected in other markers. From a taxonomy stand point, using multiple markers greatly increased the numbers of taxa detected in comparison to any single marker. Good database representation for many of the taxa detected allowed for relatively high confidence in taxonomic assignments of species known to occur in OCNMS. Due to the high sequence conservation of some genes, many closely related species have genes that are invariant in the amplified region. Invariability within amplified sequences meant many sequences with database representation could be identified to order but could not be assigned to a species, genus or even a family. This was particularly common in 18S rRNA and 16S rRNA. More complete databases would likely not resolve this problem, but other more variable amplified regions or longer sequencing technology could. Based on species accumulation curves, 44 samples was insufficient sampling for the OCNMS, and more taxa would be recovered with additional samples. However, 44 samples was enough for the detection of many of the taxa known to occur in OCNMS (Figure 1).

Figure 1) Map of Olympic Coast National Marine Sanctuary (OCNMS). Sanctuary boundary is shown in yellow, samples were collected at OCNMS seasonal moorings (purple dots), OCNMS mooring that was not sampled (white dot).

Detection alone is insufficient for comparisons between sites and years. Some level of confidence in true absence from non-detection needs to be established before managers can use this type of data to make decisions. We used occupancy models to estimate the level of occupancy for each of the ASVs and each terminal taxa for all four markers. Low probabilities of detection for ASVs/taxa were common in all markers, resulting in a wide range of confidence levels for occupancy (Figure 3). This was most prevalent in the more targeted markers (16S rRNA and 12S rRNA) where the majority of ASVs/taxa fell under probability of detection thresholds for the number of samples collected (n=5). This result is not all that surprising as these markers were specifically designed to target taxa with relatively low abundance whose sequences would likely be outcompeted by more abundant sequences during PCR amplification. The targeted nature of these primers changes the relationship between sequence reads and probability of detection. There is a strong correlation between total sequence reads of an ASV/taxa and its probability of detection with 18S rRNA and COI, but with the more targeted markers this relationship is not as well supported (Figure 3c). With few complementary binding sites, primers can over amplify compatible sequences during PCR [45]. This is a known issue with site comparisons in eDNA, and while it is possible that this over representation could occur in general markers in samples with low DNA complementation, it is more likely to happen when using targeted markers and may partially explain the difference in reads/occupancy relationship between general and specific markers. The surprising result of equivalent probabilities of detection between unicellular and multicellular organisms (Figure 3e), indicates that differences in the reads/occupancy relationship and taxa above non-detection threshold is due to samples with low initial target organism DNA.

Taxon-targeted markers do increase detection of rare sequences; both of the specific markers detected species that the general markers did not detect. Despite this detection of rare species, the targeted markers do not result in greater probabilities of detection overall. Both of the targeted markers tested had a lower proportion of high probability of detection sequences than the general markers (Figure 3c). This is not a design flaw; they are simply designed to detect relatively rarer sequences, but it does impact how eDNA data from targeted markers can be analyzed. With 20 samples the probability of detection of any given taxa needs to be over 0.14% for 95% confidence that the taxon is absent and not simply a non-detection (Table 1). At this level only 55 taxa for 16S rRNA and 14 taxa for 12S rRNA are retained compared with 179 taxa with 18S rRNA and 107 taxa with COI. As most of the taxa recovered by the specific markers fail to meet minimum levels of detection the majority of the data generated by these markers was eliminated from further analysis.

If a study’s aim is simply detection of relatively rare taxa then targeted markers may be sufficient, but if comparisons of relative abundance between sites or timepoints are important, then targeted markers may not be the best option. Only four of the eleven taxa of interest detected were above the probability of detection threshold for 20 samples. Due to the special significance of these taxa, any long-term monitoring effort in OCNMS would likely need to include these species. Rather than using targeted metabarcoding given problems due to amplification competition and probabilities of detection, it is more efficient to use species-specific markers for all key monitoring taxa either by presence-absence using simple PCR or quantitatively with qPCR or ddPCR. In addition to eliminating problems with differing amounts of template DNA, these methods are also more sensitive than amplicon sequencing, making comparisons across sites and years more reliable [46]. All three methods are compatible with occupancy models, provided sufficient replicates are collected. Because the same filter extractions can be used for metabarcoding with general markers and for species-specific quantification methods, no additional samples are required. The investment in time and cost with these additional single species assays is only slightly more than running highly targeted metabarcoding markers, and the reproducibility and ability to compare different sites and years is far greater.

Covariate testing on occupancy levels did not find significant differences in any ASVs/taxa either north/south or near shore/offshore. Based on the large proportion of ASVs/taxa low probability of detections, we had just enough samples (∼20/region) to test the most common ASVs/taxa. An NMDS analysis of sequence counts in 18S and COI suggests that the site at Makah Bay may have been distinct from other sites (data not shown), but with only 5 samples per site we did not have enough samples to test individual sites for differences using corrected occupancy levels.

Occupancy models have been employed with great success with other ecological monitoring methods such as camera traps [47], transect surveys [48], and bird call surveys [49]. They have only recently been applied to eDNA studies [38,39]. By assuming site invariance between biological replicates, occupancy models can calculate the probability of detection for individual taxa/ASVs, and from that calculated probability of detection, correct site occupancy for non-detection [42]. These calculated probabilities can be used to estimate numbers of replicates required and quantify absence versus non-detection. In addition, corrected site occupancy estimates greatly reduce the chance for type I error, allowing for greater confidence for taxa/ASVs that are significantly correlated to covariates. Designing eDNA monitoring experiments that are able to use occupancy models would significantly improve ecological monitoring projects employing eDNA.

Another benefit of using thresholds of detection calculated from occupancy models is contamination removal. Contamination is common in eDNA studies and can be difficult to remove with high confidence. In this study contaminate reads were largely at low abundance with probabilities of detection under confidence thresholds for occupancy. By only using ASV’s/taxa whose probabilities of detection were high enough for confidence in occupancy with the samples collected, all known contaminants were eliminated. This was true with all four markers. With the addition of negative controls, occupancy models may prove to be a powerful way to address contamination issues with eDNA.

Using thresholds of detection for contamination removal has several advantages to other common contamination removal methods. Thresholds of detection are based on sample number and are therefore standardized between studies and markers. One of the major drawbacks of decontamination methods which remove low-abundance ASV’s, is that the level of abundance is often arbitrary and differs between sequencing runs, studies, and markers. Another advantage with using occupancy models for decontamination is that there is no need to curate a database of “real species” thought to occur from the sampled region. The use of these curated databases are a common decontamination method and are often used to set count thresholds or even to remove anything not in the database. Custom databases create a number of issues: the database needs to be published or other researches cannot directly compare results; they may exclude taxa that are not anticipated to be present; and they may fail to distinguish false positives in the taxa that have been included in the database. Inability to distinguish contamination from “true” data is particularly problematic over multiyear monitoring projects where many taxa are expected to be sequenced repeatedly with the same markers, possibly creating significant noise in the data.

There are drawbacks from using occupancy models with eDNA. Most of these drawbacks are related to the required number of samples, a minimum of five biological replicates per site [50]. Occupancy models reduce type I error, but increase type II error particularly with insufficient sampling. Because probabilities of detection with eDNA are strongly correlated to sequence abundance (Figure 3c), it is likely that the sequence abundance and not some inherent characteristic of the taxa is influencing probability of detection. This means that probabilities of detection will need to be re-calculated every time, limiting sampling optimization. Based on the probabilities of detection we reported, 10-20 samples per site are required for monitoring OCNMS using eDNA (Figure 3, Table 1). There are strongly diminishing returns above 20 samples, and the majority of taxa/ASVs have detection probabilities below the confidence thresholds for absence with less than 10 biological replicates per site (Figure 3).

Our data indicates that from the taxonomic perspective, 44 samples were not sufficient sampling for any of the four markers tested (Figure 2b). The variation between replicates was also large enough that collection of additional samples per site is warranted, suggesting that regional monitoring studies using eDNA benefit from more intensive sampling at fewer sites. Since robust sampling is required for monitoring studies using eDNA, many of the drawbacks of occupancy models may be eliminated simply by adequate sampling.

A separate issue from sample number is that occupancy models assume that a site is invariant between sampling events. For transient and planktonic species this assumption is quickly violated in marine environments. This is not a problem when samples are collected by hand, but when samples are collected automatically from a mooring, sample collection is limited to the collection device’s capacity. High replicate sample collection will limit either sampling frequency or length of deployment forcing managers to make difficult compromises. Since the automated collection of biological time series has the potential to be a strength of eDNA, this may be a major limitation of occupancy models, however given the benefits of occupancy models in terms of standardization between sites and years, contamination removal, and robust covariate testing, the benefits may outweigh the limitations. It may be possible to incorporate occupancy models and automated sample collection, while still keeping longer time series. If the time between sampling events is short enough, it may be possible to use a rolling window of collection events that could be treated as invariant. Whether these rolling windows could be used and the length of the interval between samples will need to be tested first.

Since occupancy is based on probabilities of detection, it may be possible to decrease the number of required samples by improving probabilities of detection. There may be several ways to improve probabilities of detection, such as increasing the amount of water filtered [51], targeting key habitats for particular species [52], increasing sequencing depth [53], or selecting better markers [54]. These methods need to be tested, but five 40L water samples may be more robust than 20 1L samples. Such approaches may also be able to improve taxa/ASV detection beyond 20 samples due to the rapidly diminishing returns.

This study demonstrates how occupancy models can be used to provide a statistical framework for using eDNA for coastal community monitoring and is a step toward the ultimate goal of using eDNA to generate rich biological community data from multi-year monitoring efforts to help inform policy and management decisions.

## Methods

Samples were collected in partnership with OCNMS personnel aboard the *R/V Tatoosh* in September of 2019. Nine sites in OCNMS were selected based on the presence of long term oceanographic moorings deployed and maintained by OCNMS (Figure 1). Moorings are equipped with a variety of physical and chemical sensors (for detailed mooring schematics see (“Olympic Coast - Science - Oceanographic Moorings”)). Five one-liter water samples were collected from each of the nine sites. Samples were collected off the bow of the *R/V Tatoosh* drifting into the current to minimize potential for contamination from the hull. Sterilized one liter Nalgene bottles were rinsed three times with sea water, samples were collected at the surface and then stored in a cooler until they could be filtered within 7 hours of collection. Water samples were filtered through a 0.45µ MCE cup filter (Sterlitech, AF045W50),stored in 750 µl CTAB, and frozen at -20°C.

Filters were thawed and sterile garnet beads (Quiagen, 13123-50) were added to each tube; samples were bead beat for 15 min at RT. Following bead beating, CTAB was removed and a standard chloroform extraction was performed. All subsequent steps were done in a laminar flow hood to minimize contamination. Samples were quantified using a nanodrop (ThermoFisher) and diluted to a concentration of 10ng/µl of DNA.

Four markers were tested in this study: the V8 region of 18S rRNA as a general eukaryotic marker with good metazoan discrimination [55]; COI general marker covering half of the Barcode of Life region [56]; 16S rRNA as a metazoan specific marker [57]; and 12S rRNA targeting fish [58]. Amplification of each marker was optimized to have the fewest number of cycles (Table 2). Master mix for PCR in µl: 13.2 H_2_O, 5 5x buffer 1.5 MgCl_2_, 1 dNTPs (10 ng/µl concentration), 1 forward and reverse primer, 0.3 High fidelity Taq polymerase, 2 DNA ∼10 ng/µl concentration (New England BioLabs). Positive and negative controls were run alongside OCNMS samples. Our positive controls were genomic DNA from *Vicuna vicuna* (vicuna) 0.5 ng/µl and *Struthio camelus* (Ostrich) 0.25 ng/µl. Tissue samples for positive control species were provided by The Burke Museum UWBM: 82239 and UWBM: 91390. Negative control was from one liter RO water from our lab system. Amplification and addition of Illumina adaptors was done in a one-step PCR. Two technical replicates for each sample and marker were pooled before PCR purification. Amplified PCR products were cleaned using AMPure beads in manufacture specified concentrations (Berkman Coulter, A63881). Indexing and sequencing was done at the Oregon State’s Center for Genome Research and Biocomputing on an Illumina MiSeq with MiSeq Reagent Kit v3 2×300.

**Table 2).**
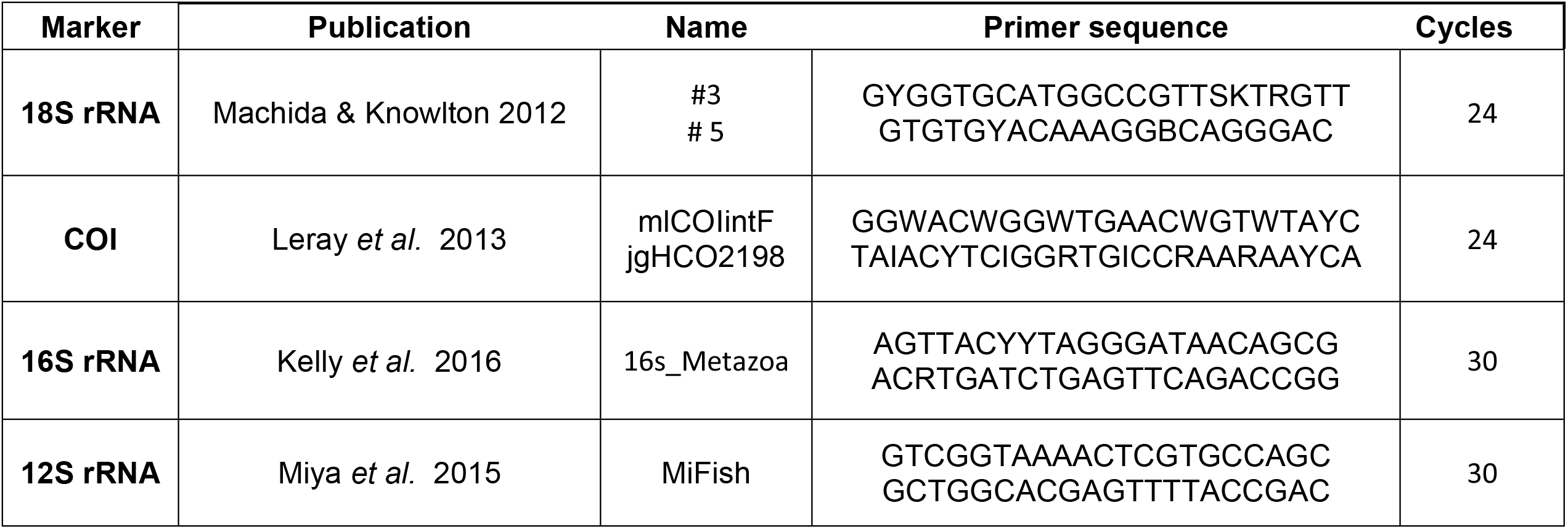
Table of markers.

Raw sequencing reads were first processed to remove primers and Illumina adaptors, trimmed with Cutadapt v 2.4 with flags *-a -A -m 100 -discard-untrimmed* [59]. Quality filtering, error correction, merging, and amplicon sequence variant (ASV) assignment was done in DADA2 v1.14 *pool=pseudo* [60]. NCBI’s BLASTn was run against the NCBI nr database. The top 50 hits were downloaded, hits were retained if they had greater than 90% coverage. Taxonomy was assigned by highest percent identity (greater than 97% to species, 95% to genus, 90% to phylum, less than 90% unknown). In cases where there was no clear best hit, taxonomy was assigned to the lowest taxonomic level possible. Sequence reads from multiple amplicon sequence variants (ASVs) with the same taxonomic assignment were merged. Unknowns, terrestrial species, and known contaminants were removed from taxonomic analysis. Basic summary statistics were done in the Vegan v 2.5-7 R package [61] and PhyloSeqv 1.16.2 R package [62].

Single species occupancy models were run for all ASVs, as well as for all assigned taxa, from all markers in the Unmarked v 1.0.1 R package [63]. Resulting probabilities of detection and occupancy estimates were used to compare the four different markers and calculate the required number of samples for sufficient statistical power. Covariate analyses were performed on three models: null (occupancy was not site dependent); near-shore (15m water depth) vs off-shore (42m water depth); and north (Teahwhit Head, Cape Alava, Makah Bay) vs south (Kalaloch, Cape Elizabeth). Models were tested for significance and Akaike information criterion (AIC) was calculated for model selection. Analysis of occupancy models and figure creation was done in R v 3.6.3 computational environment [64].

## Acknowledgements

This publication is partially funded by the Joint Institute for the Study of the Atmosphere and Ocean (JISAO) under NOAA Cooperative Agreement NA15OAR4320063, Contribution No. 2021-1148, and the Pacific Marine Environmental Laboratory (PMEL), Contribution No.XXXX. Funding for this work was provided by NOAA ‘Omics in care of Dr. Carol Stepien as well as personnel funding from the National Research Council. Special thanks to the crew of the *R/V Tatoosh*: Kathy Hough, LT. Alisha Friel, LTJG. Anna Hallingstad for their help with sample collection.

